# Perceptual responses to microstimulation frequency are spatially organized in human somatosensory cortex

**DOI:** 10.1101/2020.07.16.207506

**Authors:** Christopher L. Hughes, Sharlene N. Flesher, Jeffrey M. Weiss, Michael Boninger, Jennifer L. Collinger, Robert Gaunt

**Affiliations:** Rehab Neural Engineering Labs, University of Pittsburgh, Pittsburgh, PA; Department of Bioengineering, University of Pittsburgh, Pittsburgh, PA; Center for the Neural Basis of Cognition, University of Pittsburgh, Pittsburgh, PA; Department of Neurosurgery, Stanford University, Stanford, CA; Department of Electrical Engineering, Stanford University, Stanford, CA; Department of Physical Medicine and Rehabilitation, University of Pittsburgh, Pittsburgh, PA; Human Engineering Research Laboratories, VA Center of Excellence, Department of Veterans Affairs, Pittsburgh, PA

## Abstract

Microstimulation in the somatosensory cortex can evoke artificial tactile percepts and can be incorporated into bidirectional brain-computer interfaces (BCIs) to restore function after injury or disease. However, little is known about how stimulation parameters themselves affect perception. Here, we stimulated through microelectrode arrays implanted in the somatosensory cortex of a human participant with a cervical spinal cord injury and varied the stimulus amplitude, duration, and frequency. Increasing the amplitude and duration increased the perceived intensity on all tested electrodes. Surprisingly, we found that increasing the frequency evoked more intense percepts on some electrodes but evoked less intense percepts on most electrodes. Electrodes divided into three groups which evoked distinct perceptual qualities that depended on the stimulus frequency and were spatially organized in cortex. These results contribute to our growing understanding of the structure and function of the somatosensory cortex and will facilitate principled development of stimulation strategies for bidirectional BCIs.

## Introduction

Bidirectional brain-computer interfaces (BCI) can restore lost function to people living with damage to the brain, spine, and limbs (Collinger et al., 2018; Fetz, 2015; Flesher et al., 2019; Hughes et al., 2020). BCI users can control an end effector using neural activity recorded from motor cortex and receive sensory feedback through intracortical microstimulation (ICMS) in somatosensory cortex. Beyond the practical aim of restoring sensation to improve motor function, existing bidirectional BCIs in human participants provide an unprecedented ability to investigate the nature of sensory perception.

The behavioral effects of ICMS in somatosensory cortex have been studied in detail in non-human primates (NHPs) (Dadarlat et al., 2014; Kim et al., 2015b, 2015a; Romo et al., 2000, 1998). However, animals are limited in their ability to perform certain psychophysical tasks. NHPs can learn to discriminate between two or more stimuli, and their ability to perform these tasks can provide insight into how stimulus parameters affect sensory perception. However, they can never describe the qualitative nature of the sensory percepts they are experiencing, nor can they be trained to perform more complex psychophysical tasks such as magnitude estimation. NHP studies can therefore lead to hypotheses about how stimulus parameters affect qualitative aspects of perception, but only human studies can investigate these directly.

Limited work has been conducted in humans using ICMS of somatosensory cortex to restore sensation (Armenta Salas et al., 2018; Fifer et al., 2020; Flesher et al., 2016). From these studies we know that ICMS can evoke tactile sensations that are perceived to originate from the hands (Fifer et al., 2020; Flesher et al., 2016) and arms (Armenta Salas et al., 2018) and that the stimulation locations in the cortex that elicit these percepts agree with known cortical somatotopy (Penfield and Boldrey, 1937). Participants reported naturalistic sensations such as “pressure” and “touch” (Flesher et al., 2016) as well as “squeeze” and “tap” (Armenta Salas et al., 2018), but the quality and naturalness varied between stimulated electrodes within each participant. Additionally, all studies found that increasing the stimulus current amplitude consistently increased the perceived intensity of the tactile percepts. The effect of stimulus pulse frequency has been less studied, although low frequencies may require higher amplitudes to evoke a detectable percept (Armenta Salas et al., 2018).

It has been frequently suggested that increasing the stimulus frequency increases the perceived intensity of a stimulus train. Increasing the pulse frequency of ICMS reduced the current amplitude required to evoke a detectable percept in NHPs (Kim et al., 2015a; Romo et al., 2000, 1998) and rats (Butovas and Schwarz, 2007; Semprini et al., 2012). This was thought to indicate that increasing pulse frequency increased perceived intensity. Additionally, increasing amplitude biased NHPs (Callier et al., 2020) and rats (Fridman et al., 2010) towards selecting stimulus trains as having higher frequencies, providing further evidence that increasing pulse frequency increases perceived intensity. Perceived intensity also increases as stimulation amplitude and frequency are increased in human peripheral nerves (Graczyk et al., 2016) and human visual cortex (Schmidt et al., 1996). This is also true for mechanical stimuli where perceived intensity increased with increasing vibration frequency in able-bodied human participants using tactile input to the hand (Hollins and Roy, 1996; Muniak et al., 2007; Verrillo et al., 1969). This implies that both electrical and tactile stimulation with higher frequency components are perceived as being more intense. Our goal here was to understand if this same principle applied to ICMS of human somatosensory cortex and to evaluate whether perceptual qualities were affected by changes in stimulus pulse frequency.

In ongoing experiments, we have implanted microelectrode arrays into the motor and somatosensory cortices of a participant with a C5/C6 spinal cord injury to create a functionally useful bidirectional BCI and to study sensorimotor control in humans. With these implants, we delivered ICMS to somatosensory cortex, which evoked tactile percepts that felt like they originated from the paralyzed hand (Flesher et al., 2016). However, the percepts themselves varied considerably, from more natural sensations, such as touch and pressure, to less natural sensations, such as vibration and tingle. In order to represent more complex and intuitive tactile inputs with ICMS, it is critical that we understand how stimulus parameters directly affect sensation.

We are particularly interested in how stimulus parameters, such as current amplitude, pulse frequency, and train duration change the perceived intensity of tactile percepts. The ability to control perceived intensity in a bidirectional BCI will be essential, as modulated sensory feedback is crucial for object interaction (Johansson and Flanagan, 2009; Nowak et al., 2013). While grasp contact could be relayed by simple on-off stimulation, conveying grip force, which is essential for grasp stability, efficiency and precision (Godfrey et al., 2016; Nowak et al., 2004; Nowak and Hermsdörfer, 2006), requires the ability to modulate the perceived intensity of a stimulus. We sought to assess the effects of changing the stimulus pulse frequency on several perceptual metrics and expected to see increases in the perceived intensity as the stimulus pulse frequency increased in our participant.

## Results

### Effects of frequency on perceived intensity are electrode-dependent

We delivered ICMS trains through individual electrodes and asked the participant to report the perceived intensity on a self-selected scale, which typically ranged from 0 to 4. We found that increasing the stimulus current amplitude and train duration consistently increased the perceived intensity of the evoked sensations on all tested electrodes (Supplementary Figure 1). However, we found that the relationship between stimulus frequency and perceived intensity was electrode-dependent (Figure 1). We delivered a 60 μA stimulus train for 1 s at pulse frequencies ranging from 20 to 300 Hz. On some electrodes, percept intensity increased with stimulus pulse frequency (Figure 1B). However, on over half of the tested electrodes, the opposite effect occurred; stimulus trains with low pulse frequencies (20-100 Hz) were perceived as being the most intense and the intensity *decreased* as the stimulus pulse frequency *increased* (Figure 1C,D). Examining polynomial fits of the data, we found that electrodes could be divided into three categories: electrodes with the highest intensity response at the lowest pulse frequency (Figure 1D), electrodes with the highest intensity responses at an intermediate pulse frequency (Figure 1C), and electrodes with the highest intensity response at the highest pulse frequency (Figure 1A). Based on this observation, we used k-means clustering to separate electrodes into three categories based on the reported percept intensity at 20, 100, and 300 Hz (Supplementary Figure 2). The two most salient features that separated these groups of electrodes were the median reported intensity across all frequencies and the pulse frequency at which the maximal intensity occurred. For simplicity, we refer to these groups based on the pulse frequency range at which the maximal intensity occurred in the polynomial fit: High Frequency Preferring (HFP), Intermediate Frequency Preferring (IFP), and Low Frequency Preferring (LFP) electrodes.

**Figure 1:**
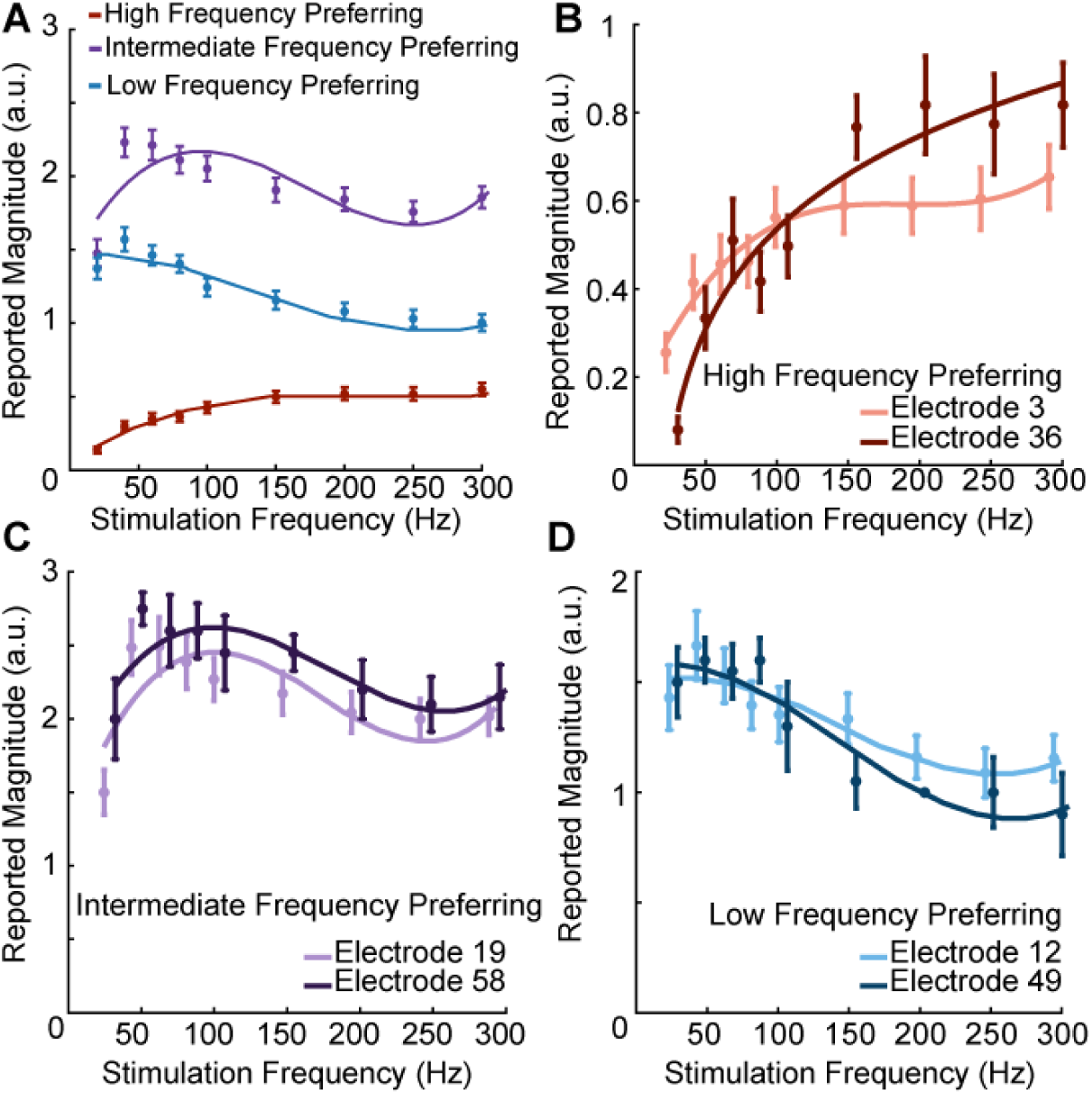
Pulse frequency drives electrode specific changes in intensity which can be grouped into three categories. A) Perceived intensity for each aggregated frequency preference group. Different colors represent different categories. Each data point shows the mean intensity response of all of the electrodes in a given category. B) Perceived intensity for two examples of high-frequency preferring electrodes that evoked the most intense percepts at the highest pulse frequencies and that generated the least intense percepts overall. C) Perceived intensity for two examples of intermediate-frequency preferring electrodes that generated the most intense overall percepts, which occurred between 40 Hz and 100 Hz. D) Perceived intensity for two examples of low-frequency preferring electrodes, which generated intermediate overall intensities that were maximized between 20 and 100 Hz. Error bars represent the standard error. Solid lines are polynomial fits of the data. Axes are scaled differently between panels for clarity.

Seven electrodes were tested in multiple sessions (3 to 6 per electrode) to determine whether the relationships between pulse frequency and perceived intensity were consistent across sessions. The perceived intensities reported on all electrodes that were tested in three or more sessions changed by statistically significant amounts as the stimulus pulse frequency changed (p < 0.001, Friedman test). Additionally, we found that the relationships between pulse frequency and normalized intensity on these electrodes – in other words, the shape of the frequency-intensity relationship – did not change significantly across test days (p > 0.05, Friedman test) (Supplementary Figure 3). An additional 22 electrodes were tested in one or two sessions. Of the 29 electrodes tested in total, 20 electrodes exhibited intensities that changed by statistically significant amounts as the stimulus frequency changed (p < 0.02, Friedman test). Of these 20 electrodes, five were classified as low-frequency preferring, seven were classified as intermediate-frequency preferring, and eight were classified as high-frequency preferring.

The pulse frequency at which the maximum intensity occurred on an electrode was significantly related to the magnitude of the reported intensities (p < 0.001, Kruskal-Wallis). Intermediate-frequency preferring electrodes had the highest median intensity of 1.94 (43% of the maximal intensity reported) and evoked the most intense percepts in the low frequency range (20-100 Hz) while high-frequency preferring electrodes had the lowest median intensity of 0.29 (6% of the maximal intensity reported) and evoked the most intense percepts in the high frequency range (150-300 Hz).

### Frequency-intensity relationships are preserved across suprathreshold amplitudes

We measured whether the frequency-intensity relationships were affected by stimulus current amplitude. If the frequency-intensity relationships were dependent on the current amplitude, this result might reflect idiosyncratic recruitment effects of ICMS. Therefore, we presented stimulus trains at three current amplitudes (20, 50, and 80 µA) and three pulse frequencies (20, 100, and 300 Hz), which spanned the range of detectable and safe parameters, and asked the participant to report the perceived intensity of the evoked percepts. There were no significant differences in the shape of the frequency-intensity relationships for the three electrode groups at 50 and 80 μA after controlling for changes in median intensity caused by increasing current amplitude (p = 0.21-0.99 Friedman’s test, Figure 2). The reported intensity on low-frequency preferring electrodes peaked at 20 Hz at both current amplitudes (p = 0.02, Kruskal-Wallis, Figure 2A), whereas the reported intensities of intermediate-frequency preferring electrodes peaked at 100 Hz for both current amplitudes (p < 0.001, Kruskal-Wallis, Figure 2B) and the reported intensity on high-frequency preferring electrodes peaked at 300 Hz for both current amplitudes (p < 0.001, Kruskal-Wallis, Figure 2C). Interestingly, when we decreased the current amplitude to 20 μA, which was close to the detection threshold for most electrodes, increasing the pulse frequency from 20 to 100 Hz evoked more intense percepts for all electrode groups (p < 0.05, Kruskal-Wallis, Figure 2A,B,C, 20 µA). There were highly significant differences between the shape of the frequency-intensity relationships for all groups at 20 μA versus 50 or 80 μA (p < 0.001, Friedman’s test) even after controlling for changes in the median intensity caused by increasing current amplitude. At 20 μA, the percept intensity was very low, making magnitude estimation akin to a detection task.

**Figure 2:**
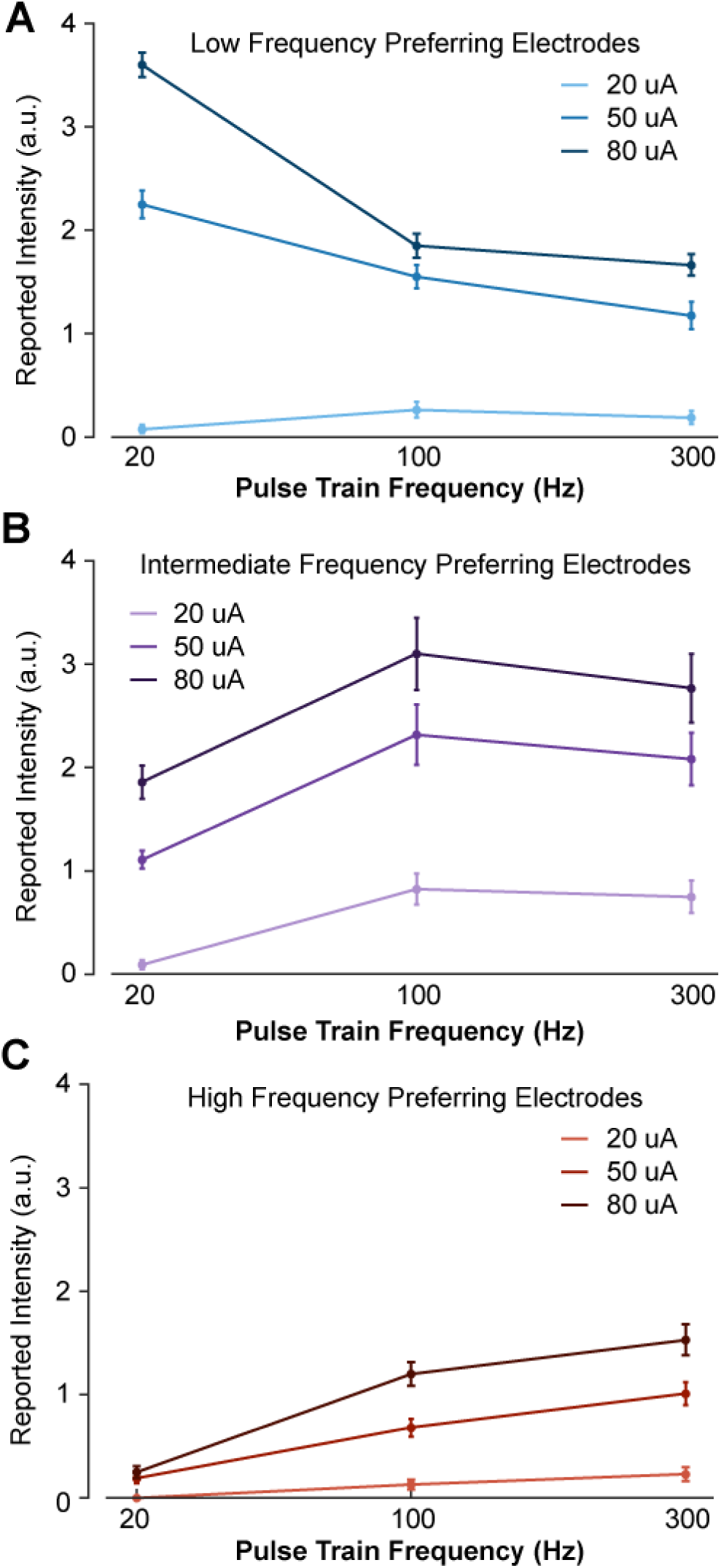
Stimulus current amplitude does not change the relationship between pulse frequency and intensity at suprathreshold amplitudes. Magnitude estimation data for different current amplitudes and pulse frequencies. Data were aggregated across electrodes by their category, where each plot shows a different category of electrodes. Perceived intensity values for A) LFP electrodes, B) IFP electrodes, and C) HFP electrodes at different current amplitudes and pulse frequencies. Different colored bars represent different current amplitudes. Error bars indicate the standard error across electrodes. We tested two LFP electrodes, three IFP electrodes, and two HFP electrodes which were each tested twice in different sessions.

### High frequency stimuli are detected more reliably at perithreshold amplitudes

Our observation that higher stimulus pulse frequencies can evoke less intense percepts at suprathreshold stimulus current amplitudes seems to contradict observations that higher frequencies can evoke detectable percepts at lower amplitudes in NHPs (Kim et al., 2015a; Romo et al., 2000, 1998). These detection threshold experiments suggest that increasing the stimulus frequency should generally evoke more intense percepts. However, the effect of changing ICMS parameters on perceived intensity cannot be tested directly in NHPs. Indeed, we found that the perceived intensity at the lowest tested currents always increased when the frequency increased from 20 to 100 Hz (Figure 2A,B,C, 20 µA), but that this effect was not always maintained at higher current amplitudes (Figure 2A,B, 50 and 80 µA). To explicitly compare our results to NHP work, we performed a detection task in which the current amplitude was set to perithreshold levels and the pulse frequency was varied between 20, 100, and 300 Hz. We found that at 300 Hz, the interval containing the stimulus train was correctly identified 80% of the time across all tested electrodes (Supplementary Figure 4). Similarly, when the pulse frequency was set to 100 Hz, the mean detection accuracy was 72%. In contrast, when the pulse frequency was set to 20 Hz, the mean detection accuracy was just 42%, which was below chance levels of 50%. Detection accuracies at 100 Hz and 300 Hz were significantly higher than the detection accuracy at 20 Hz (p < 0.05, ANOVA) but were not significantly different than each other (p = 0.66, ANOVA). These results are consistent with the observations in NHPs but demonstrate a dichotomy between detection and intensity evaluation: detection thresholds are not necessarily predictive of perceived intensity and predictions about subjective measures of perception may be difficult to make in animals.

### Frequency-intensity relationships are associated with different perceptual qualities

One advantage of studying somatosensation in humans is the ability to document the sensory qualities evoked by stimulation (Table 1). We found that there were significant differences in the qualities evoked on electrodes belonging to different categories defined by the effect of pulse frequency on intensity (Figure 3A). Additionally, the sensory qualities for electrodes in each group were differentially modulated by pulse frequency (Figure 3B).

**Table 1:** All reported percepts and their percent occurrence at each pulse frequency. Since multiple percepts can be reported for a single stimulus, columns will add to more than 100 percent. There were 152 samples at 20 Hz, 621 samples at 100 Hz, and 85 samples at 300 Hz.

|  |  | % of reports that contained descriptors |  |  |
| --- | --- | --- | --- | --- |
| Descriptors |  | 20 Hz | 100 Hz | 300 Hz |
| Descriptors included from the start | Tingle | 26 | 67 | 72 |
|  | Pressure | 50 | 35 | 31 |
|  | Warm | 7 | 31 | 31 |
|  | Sharp | 7 | 12 | 11 |
|  | Vibration | 1 | 5 | 8 |
|  | Touch | 26 | 1 | 1 |
| Descriptors developed by participant | Buzzing | 2 | 9 | 9 |
|  | Tapping | 32 | 1 | 2 |
|  | Sparkle | 16 | 0 | 0 |
|  | Prick | 3 | 4 | 2 |

**Figure 3:**
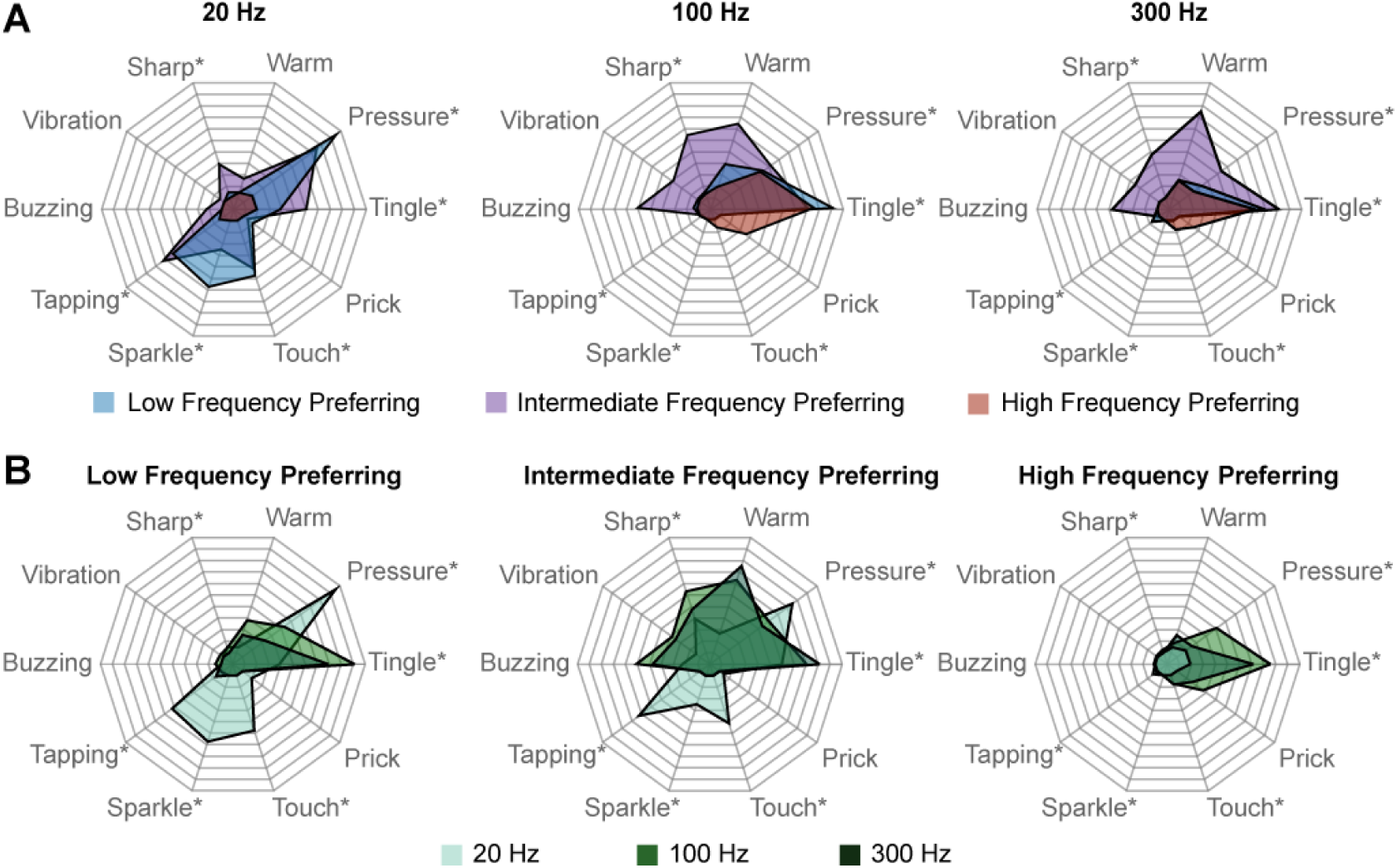
Perceptual qualities are associated with specific electrode categories and stimulus pulse frequencies. Radar plots showing the distribution of reported qualities at different pulse frequencies for each electrode category. A) Percepts sorted by pulse frequency. Electrode categories are indicated with different colors. B) Percepts sorted by electrode categories. Pulse frequencies are indicated with different colors. In each plot, qualities on which there was a significant difference between categories, as determined with Fisher’s exact test, are marked with an asterisk.

At 20 Hz, we found that LFP and IFP electrodes evoked percepts with pressure, tapping, sparkle, and touch qualities. These qualities were not evoked on HFP electrodes at any frequency. At this low stimulation frequency, HFP electrodes were generally not detectable, resulting in few reports of any percepts. At 100 Hz, IFP electrodes evoked percepts with buzzing, vibration, and sharp qualities. LFP and HFP electrodes never evoked these qualities when stimulated at 100 Hz. HFP electrodes also evoked sensations of touch and prick at 100 Hz that never occurred on LFP or IFP electrodes at any frequency. However, these qualities occurred on less than 30% of trials on HFP electrodes. At 300 Hz, the responses were similar to those at 100 Hz except that all electrode categories evoked less pressure.

We also clustered electrodes based on the verbal reports of percept quality at all frequencies. Interestingly, these clusters were remarkably similar to those based on intensity responses at different frequencies (Supplementary Figure 5). That these electrode categories were nearly identical when created using completely different data sets–perceptual qualities and perceived intensities–strongly suggests that these two features are measures of the same underlying properties of the neurons recruited by stimulation.

### Perceptual responses are spatially organized in cortex

Finally, we asked whether the categorization of an electrode, which corresponds to its frequency-intensity responses and evoked perceptual qualities, was related to its location in cortex. We compared the observed spatial occurrence of electrode category with a simulation that randomly assigned each category to one of the tested electrode locations while maintaining the same number of electrodes in each category. Using the simulation, we found that there was significant clustering of electrodes in the same category (Figure 4) on both the lateral array (pseudo-p = 0.003, LISA) and the medial array (pseudo-p = 0.02, LISA). This was particularly apparent on the lateral array.

**Figure 4:**
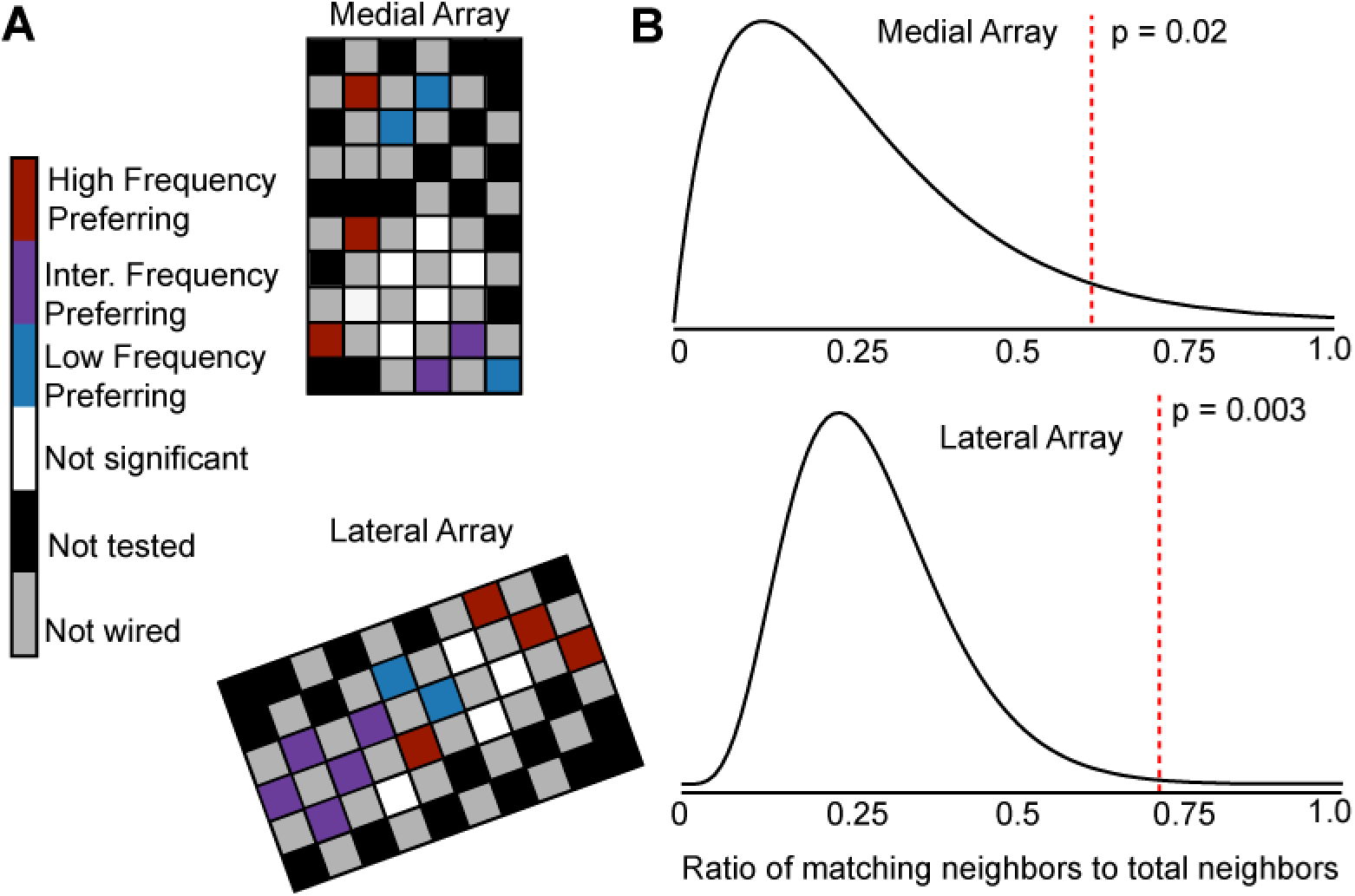
Electrode location is significantly related to electrode categorization. A) Map of the medial electrode array (top) and lateral electrode array (bottom) implanted in somatosensory cortex and the distribution of the frequency preference categorizations. The electrode arrays were implanted close to the central sulcus with the left edge of the medial array being approximately parallel to the central sulcus. The arrays are oriented to reflect the implant orientation. Colored squares represent different types of electrodes as indicated by the color bar. B) The distribution of clustering coefficients obtained from randomly distributing the medial array (top) and lateral array (bottom) electrode categorizations on a simulated array. The black curve shows a gamma distribution fit to the simulation output. The red dotted line indicates the actual clustering coefficient obtained from real data.

The clusters ranged in size from about 1-2.5 mm. However, our sampling of the cortical surface was sparse, making it difficult to assess the true size of the clusters. Furthermore, we observed that in some cases, clusters of electrodes from different categories evoked percepts from the same region of the hand. For example, LFP, IFP, and HFP electrodes elicited sensations on the palmar region beneath the middle and ring fingers. As a result, percepts from a single region of the hand, such as the index finger, could be evoked by electrodes that generated multiple response types.

## Discussion

### Neural populations preferentially respond to different stimulus frequencies in somatosensory cortex

We found that the pulse frequency of ICMS alters the perceived intensity (Figure 1) and quality (Figure 3) in an electrode-specific manner. Further, we found that electrodes with similar intensity responses and evoked qualities clustered spatially in somatosensory cortex (Figure 4). This strongly implies that the observed electrode-specific relationships between frequency and perception are not caused by random factors and are instead related to the underlying structure and function of the cortex.

Studies in somatosensory cortex using both electrophysiology (Mountcastle et al., 1969; Sur et al., 1984, 1981) and optical imaging (Chen et al., 2001; Friedman et al., 2004) techniques have demonstrated that different neural populations are sensitive to tactile input with specific frequency content, and that these populations are organized spatially in the cortex. Populations of cortical neurons were classified based on the type of peripheral mechanoreceptors that had similar responses to the same frequency ranges (rapidly adapting, slowly adapting, and Pacinian corpuscle type). However, most cortical neurons have heterogeneous input from different classes of mechanoreceptors and describing the cortical responses as being driven only by mechanoreceptors of a particular sub-modality is misleading (Pei et al., 2009; Reed et al., 2010; Saal and Bensmaia, 2014). If cortical activity simply reflected the convergent input of different mechanoreceptor sub-modalities, we would expect that the perceptual response to electrical stimulation would be uniform given the same stimulus input. In fact, experiments in NHPs showed that ICMS pulse frequency discrimination was dependent on the stimulated population and its associated modality (Romo et al., 2000). Similarly, our results demonstrate that different populations of neurons within somatosensory cortex respond preferentially to different stimulation pulse frequencies. Preferential frequency responses in cortex then are not simply driven by the response properties of an associated mechanoreceptor; rather, the somatosensory cortex is organized to preferentially encode stimulus frequencies and associated qualities.

Our data support the idea that the frequency content of a mechanical or electrical input is encoded in the somatosensory cortex and divided into patches within the overlying somatotopy. Although this idea that the cortex is organized for preferential encoding of frequency content deviates from the proposal that the cortex is tuned to specific mechanoreceptors made in previous studies (Chen et al., 2001; Friedman et al., 2004; Mountcastle et al., 1969; Sur et al., 1984, 1981), the idea that somatosensory neurons act as spatiotemporal filters to encode particular stimulus features is not entirely new (Dicarlo et al., 1998; Saal and Bensmaia, 2014). This organization in somatosensory cortex then results in different preferential responses to ICMS frequency and different evoked qualities.

### Possible mechanisms for heterogeneous perceptual responses to stimulus frequencies in cortex

The observed effects in our study must be related to different cellular responses to stimulation in different regions of the cortex. In fact, different stimulation frequencies in mouse somatosensory cortex can alter the activation of distal neurons (Michelson et al., 2019). Specifically, high pulse frequencies lead to fast habituation of activated distal neurons while low pulse frequencies can maintain this activation. Decreases in distal neuron activity at high frequencies could potentially drive decreases in perceived intensity. If neural populations differed in their ability to drive distal populations, this could result in population-dependent frequency responses.

A potential mechanism to explain electrode-dependent responses are varying distributions of inhibitory and excitatory neurons. The presence of more inhibitory neurons in a local region could result in stronger inhibitory drive at higher frequencies, resulting in more robust responses to low frequency stimuli. Indeed, recruitment of inhibitory Martinotti cells in the somatosensory cortex of rats increases as the duration and frequency of presynaptic action potentials increase (Kapfer et al., 2007; Silberberg and Markram, 2007). Further, rostrocaudal heterogeneity of inhibition has been documented in rat olfactory cortex (Large et al., 2018; Luna and Pettit, 2010). Whether such organization exists in human somatosensory cortex remains to be seen.

Short term plasticity (Tsodyks and Markram, 1997) at synapses driven by stimulation may also explain the observed effects. If a synapse is unable to resupply neurotransmitter at a rate faster than the stimulus frequency, transmitter release at the synapse could become depressed. In this scenario, neurons would be unable to recruit other neurons in synchrony with stimulation, which could result in lower recruitment and lower perceived intensity. If cells in somatosensory cortex have different time constants for transmitter recovery, this could serve as a mechanism for frequency filtering (Rosenbaum et al., 2012). Elucidating the precise mechanisms underlying observed frequency responses in cortex will require further studies in animal models.

### Human perception of ICMS contradicts predictions from non-human primate studies

Higher stimulus pulse frequencies decreased the current amplitude required to detect a stimulus train in NHPs (Kim et al., 2015a). It was predicted then that higher stimulus frequencies would increase the perceived intensity of a stimulus train. Similar to these animal studies, we found that higher frequencies improved the detectability of stimulus trains at perithreshold amplitudes. However, at suprathreshold current amplitudes, increasing the frequency did not always produce higher perceived intensities. Detection and perceived intensity then may be related, but our results suggest that they are distinct measures of perception.

In other NHP experiments, animals were trained to identify which of two intervals contained the higher frequency stimulus train, regardless of current amplitude, to determine if changes in frequency could be perceived independently of changes in amplitude (Callier et al., 2020). Increasing the amplitude always biased the animals towards selecting a stimulus train as having a higher frequency. However, animals were still able to distinguish between changes in amplitude and frequency on some electrodes. In our experiments, LFP and IFP electrodes, which generated high intensity percepts at low frequencies, often evoked percepts with highly salient qualities, such as tapping or buzzing. The presence of these qualities at some frequencies and not others (Figure 3) would allow the participant to distinguish between increases in amplitude, which only increases the percept intensity (Supplementary Figure 1), and increases in frequency, which changes the percept quality and intensity (Figure 3). On electrodes without highly salient frequency-dependent qualities, such as the high-frequency preferring electrodes, it would be difficult to disambiguate changes in amplitude and frequency.

However, an important difference between our study and the NHP studies is that many electrodes in our study evoked less intense percepts as the pulse frequency increased, which was not observed in NHPs. The reason for this is unclear, and it may be related to the larger frequency range explored in this study or even the electrode location in the cortex. However, animals cannot directly report the perceived intensity of a set of stimuli on an open scale, as is commonly done in humans. Rather, perceived intensity, as well as other subjective aspects of perception such as quality and naturalness, must be inferred from other experimental paradigms, which makes it difficult to assess how ICMS affects subjective aspects of perception in animals. This demonstrates that human experiments are crucial to understanding how ICMS modulates tactile perception, particularly for subjective evaluation of experience.

### Limitations of study

Although conducting studies in humans yields new understanding of how stimulation affects perception, there are limitations. First, these experiments were conducted in a single participant with a chronic implant. Different participants, with different timelines of injury preceding implant, could potentially respond differently to stimulation, particularly if the electrodes are implanted in a different part of the somatosensory cortex. However, the repeatability of our findings suggest that these effects were at least not due to day-to-day variations. Our participant also had limited residual sensation in his hand, which made it difficult to measure responses in cortex to tactile indentation. Comparing perceptual responses to ICMS with cortical responses to tactile indentation could help better relate these findings to previous studies in monkeys. Another potential confound is that perceived intensity can change throughout a session. Because we used pseudo-randomized presentation of different stimulus parameters to ensure that electrodes were not tested in the same order and excluded the first block of trials from analysis for each set for magnitude estimation, we believe that this phenomenon did not affect our results. We additionally cannot know if electrodes across the array are in different layers of cortex. Different layers of cortex may drive different perceptual responses with the same input. However, if this were the case, this would still reflect important functional differences in cortex which need to be understood for bidirectional BCIs. Finally, a challenge for developing mechanistic explanations of these observations is that there are few neuroscientific tools that we can use to further probe these effects in a human. Because of this, addressing the neurophysiological mechanisms of these frequency responses is difficult in a human participant and further investigation of these properties in animal models is needed.

### Implications for prostheses

Knowing that different electrodes in cortex have different perceptual properties based on the recruited population can inform our approach to creating a functional bidirectional BCI in two primary ways. First, these results may help identify electrodes that have perceptual or intensive properties that are relevant to the task being performed. Certain patches of cortex are more involved in representing specific perceptual qualities, and these patches could be selectively activated according to the tactile input to the prosthetic device. Second, these results suggest that electrode-specific stimulation encoding schemes would be particularly useful. For example, electrodes that represent “tapping” sensations could receive large amplitude transients, while electrodes that do not evoke this sensation could receive low amplitude, sustained stimulation. These methods, in conjunction with other biomimetically informed stimulus encoding algorithms, could improve the naturalness and usefulness of somatosensory feedback, in turn improving the performance of bidirectional BCIs and ultimately improving the quality of life for people living with paralysis.

## Acknowledgements

We would like to acknowledge N. Copeland for his extraordinary commitment to this study and insightful discussions with the study team, as well as Debbie Harrington (Physical Medicine and Rehabilitation) for regulatory management of the study. This work was supported by the Defense Advanced Research Projects Agency (DARPA) and Space and Naval Warfare Systems Center Pacific (SSC Pacific) under Contract N66001-16-C4051 and the National Institute of Neurological Disorders and Stroke of the National Institutes of Health under Award Numbers UH3NS107714 and U01NS108922. SNF was supported by an NSF Graduate Research Fellowship under grant number DGE-1247842. Any opinions, findings and conclusions or recommendations expressed here are those of the authors and do not necessarily reflect the views of DARPA, SSC Pacific, or the National Institutes of Health. The funders had no role in the study design, data collection, interpretation of the results, or the decision to submit this work for publication.

## Author Contributions

CLH, SNF, and RAG designed the study. CLH, SNF, and JED conducted the experiments, and built tools to collect the data. CLH analyzed the data. CLH and SNF produced the figures. MLB, JLC and RAG provided oversight and guidance for the study and participated in data interpretation. MLB is the sponsor-investigator of the study and was responsible for the participants ongoing clinical care. CLH wrote the manuscript with RAG and all authors reviewed, edited, and approved the final manuscript.

## Methods

### Participant and Implants

This study was conducted under an Investigational Device Exemption from the U.S. Food and Drug administration, approved by the Institutional Review Boards at the University of Pittsburgh (Pittsburgh, PA) and the Space and Naval Warfare Systems Center Pacific (San Diego, CA), and registered at ClinicalTrials.gov (NCT0189-4802). Informed consent was obtained before any study procedures were conducted. The purpose of this trial is to collect preliminary safety information and demonstrate that intracortical electrode arrays can be used by people with tetraplegia to both control external devices and generate tactile percepts from the paralyzed limbs; this manuscript presents the analysis of data that were collected during the participants involvement in the trial, but does not report clinical trial outcomes.

A single subject was implanted with two microelectrode arrays (Blackrock Microsystems, Salt Lake City, UT) in the somatosensory cortex. Each electrode array consisted of 32 wired electrodes arranged on a 6×10 grid with a 400 μm interelectrode space, resulting in a device with an overall footprint of 2.4 x 4 mm. The remaining 28 electrodes were not wired due to technical constraints related to the total available number of electrical contacts on the percutaneous connector. Electrode tips were coated with a sputtered iridium oxide film (SIROF). Additional details on these implants have been published elsewhere (Flesher et al., 2016).

### Stimulation protocol

Stimulation was delivered using a CereStim C96 multichannel microstimulation system (Blackrock Microsystems, Salt Lake City, UT). Pulse trains consisted of cathodal phase first, current-controlled, charge-balanced pulses which could be delivered at frequencies from 20-300 Hz and at amplitudes from 2-100 μA. The cathodal phase was 200 μs long, the anodal phase was 400 μs long, and the anodal phase was set to half the amplitude of the cathodal phase. The phases were separated by a 100 μs interphase period. At the beginning of each test session involving stimulation, we sequentially stimulated each electrode first at 10 μA and 100 Hz for 0.5 s and then at 20 μA and 100 Hz for 0.5 s. During these trials, the interphase voltage on each electrode was measured at the end of the interphase period, immediately before the anodal phase (Cogan, 2008). If an electrode’s measured interphase voltage was less than -1.5 V, the electrode was disabled for the day (Flesher et al., 2016). This step was performed to minimize stimulation on electrodes that might potentially experience high voltages, which could result in irreversible damage.

### Magnitude Estimation

We assessed the effect of stimulus parameters on perceived intensity using a magnitude estimation task. To test the potential effect of pulse frequency on intensity, pulse trains were delivered for 1 s at 60 µA with frequencies of 20, 40, 60, 80, 100, 150, 200, 250 and 300 Hz. Following each pulse train, the participant was asked to report the magnitude of the perceived intensity on a self-selected scale. The participant was instructed to use values such that a value twice as large as a previous value was twice as intense, and a value half as large was half as intense. These values typically ranged from zero to four. Each set of stimulus pulse frequencies was presented six times, with the presentation order randomized in each block. The responses from the first block were not used in the analysis to allow the participant to establish a baseline for reporting for the session. Data collected on the same electrode over multiple sessions were aggregated for analysis. Data were fit using a 1^st^ to 3^rd^ degree polynomial for visualization. Polynomial order was determined using the corrected Akaike information criterion (AICc).

We also assessed the effect of changing the stimulus current amplitude on perceived intensity while the stimulus pulse frequency was held constant. The pulse frequency was set to 100 Hz, the train duration to 1 s, and the current amplitude ranged from 20 to 80 μA in 10 μA increments. Data were fit with a linear function. Finally, we assessed the effect of changing the stimulus train duration on perceived intensity. The stimulus pulse frequency and current amplitude were set to 100 Hz and 60 μA, respectively, and the train duration was set to 0.1, 0.2, 0.3, 0.4, 0.5, 0.75, 1, 1.5, and 2 s. Data were fit with a logistic function. For current amplitude and train duration plots, the data were normalized to the median intensities of the set in which it was collected for visualization purposes.

To investigate the interaction between current amplitude and pulse frequency, we additionally tested frequency and amplitude pairs. The train duration was set to 1 s, the current amplitude was set to 20, 50, or 80 μA, and the pulse frequency was set to 20, 100, or 300 Hz. All frequency and amplitude combinations were tested for each tested electrode six times, and the first trial was excluded from analysis. Each tested electrode was tested twice on two different test sessions, resulting in ten total trials for each frequency and amplitude pair. For analysis and plotting, we divided electrodes into the categories defined in the frequency magnitude estimation described previously. We tested two LFP electrodes, two IFP electrodes, and two HFP electrodes.

### K-means clustering

Electrodes were divided into three categories using k-means clustering using the reported intensity at 20, 100, and 300 Hz. Both silhouette and elbow analysis were used to validate that k = 3 was a suitable parameter choice. We labeled the categories as Low Frequency Preference (LFP), Intermediate Frequency Preference (IFP), and High Frequency Preference (HFP) based on the frequency at which the maximum intensity occurred for the polynomial fit.

Electrodes were additionally clustered based on the reported perceptual qualities at 20, 100 and 300 Hz. Each reported quality (of which there were 10) was summed across sessions and pulse frequencies for each electrode. The total number of reports for each quality was then divided by the maximum number of reports for any electrode, so that each quality was represented by number between zero and one and contributed equally to the clustering of each electrode. No dimensionality reduction was used and electrodes were clustered within the 10 dimensions of reported qualities.

### Detection Thresholds

Detection thresholds were determined using a two-alternative forced choice task. The participant was instructed to focus on a fixation cross on a screen located in front of him. Two 1-s-long windows, separated by a variable delay period, which averaged 1-s in length, were presented and indicated by a change in the color of the fixation cross. Stimulation was randomly assigned to one of the two windows. After the last window, the fixation cross disappeared, and the participant was asked to report which window contained the stimulus.

A one-up three-down staircase method was used so that if the participant correctly identified the window containing the stimulus in three consecutive trials, the current amplitude was decreased for the next trial (Leek, 2001; Levitt, 1971). If the participant incorrectly identified the window containing the stimulus, the current amplitude was increased for the next trial. The current amplitude started at 10 μA and was increased or decreased by a factor of 2 dB. The pulse frequency was held constant at 100 Hz. This method reduced the time spent testing uninformative values but does not guarantee that all current amplitudes will be tested the same number of times. After five changes in the direction of the stimulus current amplitude (increasing to decreasing, or decreasing to increasing), the trial was stopped. The detection threshold was calculated as the average of the last 10 values tested before the fifth direction change.

We also conducted standard detection trials where the stimulus pulse frequency was changed while the stimulus current amplitude was held constant. The current amplitude was set to 1.2X the detection threshold for each electrode measured at 100 Hz. The tested frequencies were 20, 100, and 300 Hz and each pulse frequency was presented 30 times. Pulse frequencies were interleaved randomly resulting in 90 trials per tested electrode.

### Surveys

Surveys were conducted once every month from the time the arrays were implanted. During a survey, each enabled electrode was stimulated sequentially using a 1-s pulse train at 60 μA. These parameters were selected because they were typically able to evoke sensations consistently while remaining well below our maximum stimulus current amplitude of 100 μA. Typically surveys were conducted once a month at 100 Hz, but we collected additional surveys at 20 and 300 Hz. This resulted in 152 samples at 20 Hz, 621 samples at 100 Hz, and 85 samples at 300 Hz. Surveys were conducted to quantify stimulus-evoked tactile percepts. No visual or auditory cue was provided to the participant to indicate when stimulation was occurring. The participant was instructed to indicate when a sensation was detected, at which point progression through the trial was paused. The participant verbally reported when he detected a sensation, and the pulse train was repeated as many times as necessary for the participant to be able to accurately describe the location and quality of the sensation. A drawing of the hand was partitioned into different segments and the participant reported on which segments the sensation was felt. The participant also used a tablet and stylus to circumscribe the precise areas where sensation was felt on a map of the hand.

After the location of the percept was established, the participant reported the quality of the sensation using the descriptors in Table 1. The participant’s response was documented by the experimenter and video recordings were also taken during all responses. If the participant felt that the sensation was not accurately described by the provided descriptors, his response was recorded, and the best approximation using the descriptors was used. The descriptors included a five-point scale for naturalness ranging from totally unnatural to totally natural, the location of the sensation on or below the skin surface, and an assessment of pain ranging from 0 to 10. The quality of the sensation was further assessed using the following descriptors: mechanical (touch, pressure, or sharp), movement (vibration or movement across the skin), temperature (warm or cool), and tingle (electrical, tickle, or itch). These descriptors were based on a previously described questionnaire (Heming et al., 2010). The participant could report multiple qualities for a single stimulus, and in some cases, the subcategories (for example, electrical, tickle, or itch) could not be described. The participant also reported qualities that deviated from the descriptors. The participant developed four new descriptors that were not originally included, which often were combinations of the other descriptors. We attempted to reidentify these percepts in the context of a new questionnaire which was published during this study in consultation with the participant (Kim et al., 2018). Three of these sensations were reidentified as “tapping,” “buzzing,” and “prick”.

One descriptor the participant reported, “sparkle,” could not be reidentified with the new questionnaire. The participant described this percept as feeling like tapping that varied in intensity and moved around the projected field in a random manner. It should be noted that all percepts in our study were identified as tactile percepts and no proprioceptive sensations were evoked.

The survey data included in these analyses were collected during the same year as the frequency magnitude estimation data to ensure the evoked sensations were consistent across paradigms, which included data from post-implant days 630 to 962.

### Statistics

For all statistical tests, we determined whether to use parametric or non-parametric tests based on the normality of the data as assessed with an Anderson-Darling test. If the data were significantly different than normal, then we used non-parametric tests. Any time multiple comparisons were made, we used the Benjamini-Hochberg procedure to correct for multiple comparisons which resulted in a critical p-value that was used as a cut-off. If no values were significant, then the critical p-value returned is 0 and not reported and no values are considered significant. For any tests that required post-hoc comparisons, we used Tukey’s HSD test.

For magnitude estimation data, we used Friedman’s test to assess significant differences between the intensity responses at different pulse frequencies as well as differences between electrode responses across days. Friedman’s test also allowed us to compare significant effects of pulse frequency on intensity across multiple sessions by excluding experimental day as a cofactor. When comparing the same electrode across sessions, we compared intensity responses with the same tested pulse frequency and corrected for multiple comparisons. We compared differences in the median intensity of electrodes within each category using a Kruskal-Wallis test.

For detection data, we used an ANOVA to assess significant differences in the detection accuracy at different pulse frequencies.

For quality data obtained from surveys, we used Fisher’s exact test to evaluate if there was a relationship between the categorization of each electrode and the perceptual qualities evoked on the electrode. Contingency tables were developed for each descriptor and responses were row-divided by the three categories (LFP, IFP, and HFP) and column-divided by the presence or absence of the quality. Each category was compared pairwise. Fisher’s exact test was used instead of a chi-squared because the sample sizes for each group were relatively small.

To test if there was spatial clustering of the effects of frequency on perceived intensity across the array, we adopted a technique used in geographic information systems, where they are described as Local Indicators of Spatial Association (LISA) (Anselin, 1995). We quantified the number of electrodes that had an adjacent electrode with the same frequency response category and divided this by the total number of adjacent electrodes to obtain a fraction. We then randomly distributed the categorized electrodes on a simulated array with the same tested electrode locations. We conducted this simulation 100,000 times and compared the output values of this random simulation to the observed values. A pseudo p-value was obtained by comparing the total number of simulations that had a fraction greater than or equal to the observed fraction, which indicates the probability of obtaining our observed value by chance.

For all statistics, we considered p < 0.05 to be significant, and p < 0.001 to be highly significant.

---

**Supplementary Figure 1:**
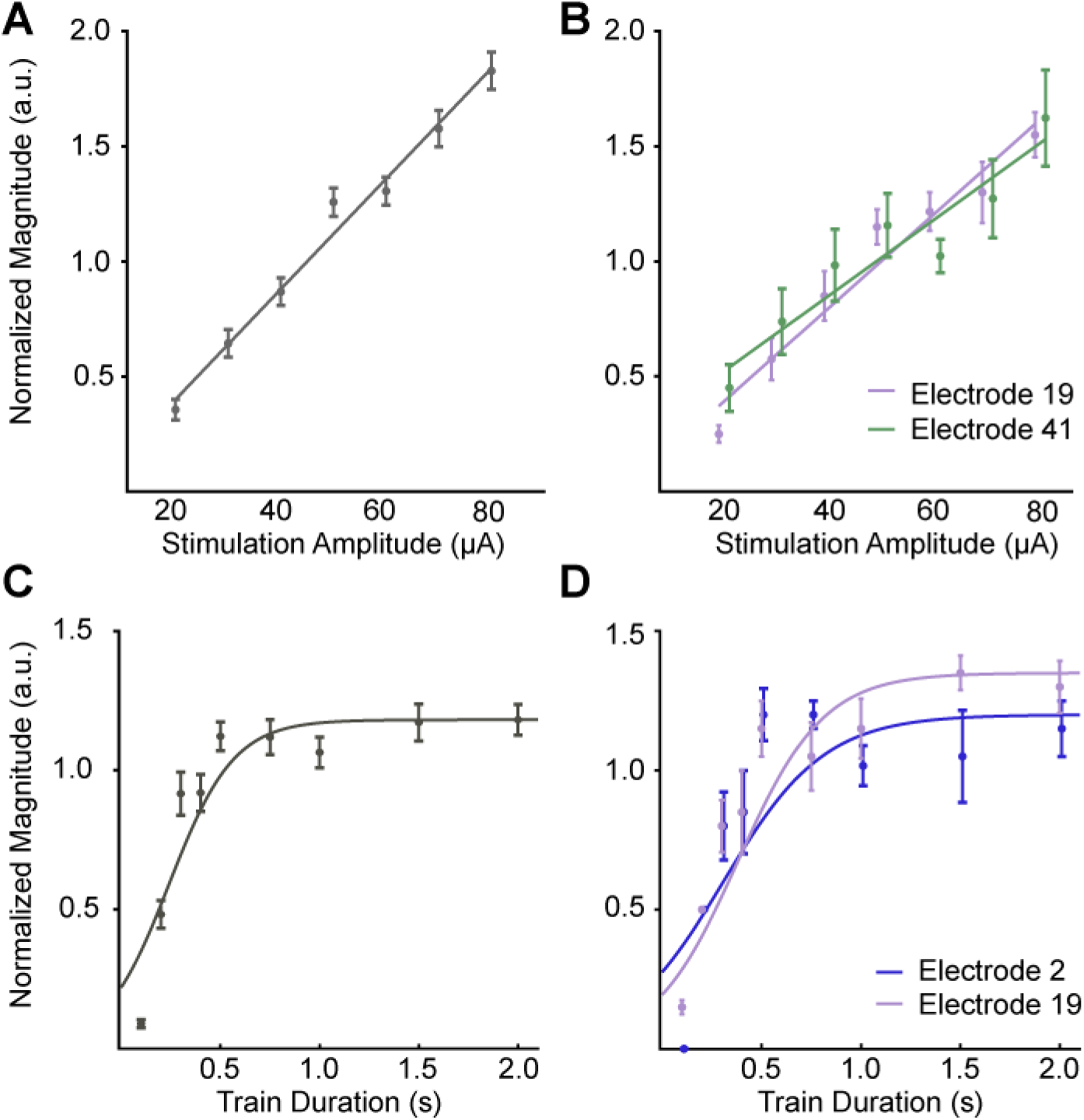
Increases in current amplitude and train duration consistently drive increases in perceived intensity. A-B) Normalized intensity as a function of current amplitude for all nine tested electrodes (A), and for two individual electrodes (B). The data were fit with a linear function. C-D) Normalized intensity as a function of train duration for all four tested electrodes (C) and two individual electrodes (D). The data were fit with a logistic function. In all panels, data points are the median reported intensity at each stimulus parameter. Samples were normalized to the median intensity value for each test. Error bars show the standard error. Note that the Y-axes are scaled differently for each panel for clarity. Colors represent different electrodes as indicated by the legends. Data points for individual electrodes are jittered slightly on the x-axis for visualization.

**Supplementary Figure 2:**
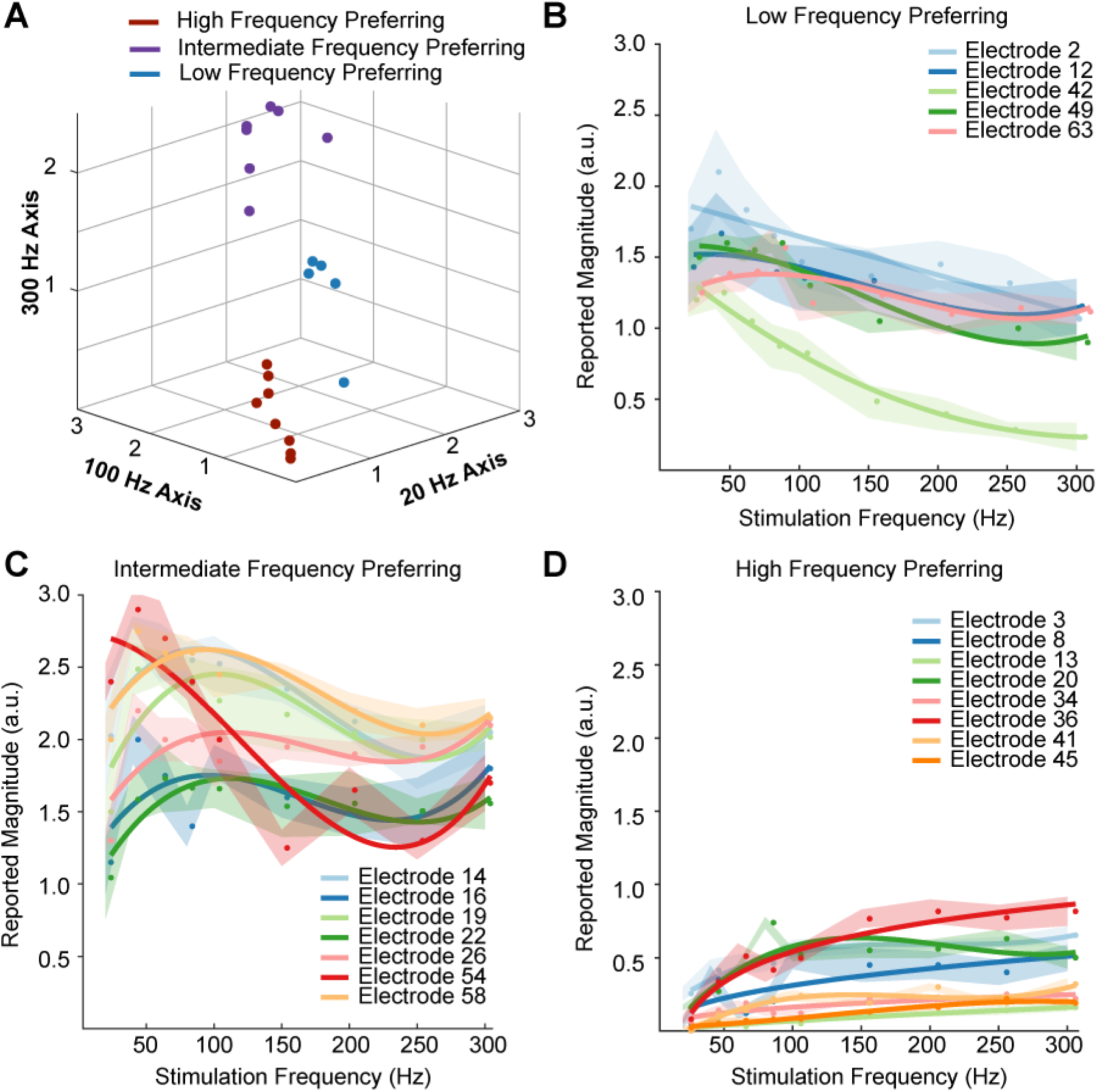
Electrodes divide into three categories based on their frequency-intensity relationships. A) K-means clustering of individual electrodes based on intensity responses at 20, 100, and 300 Hz. Individual data points are the median intensities at each frequency across all repetitions. B-D) Perceived intensity responses at different frequencies for all electrodes classified as low-frequency preferring (B), intermediate-frequency preferring (C), and high-frequency preferring (D). Shaded regions show the smoothed standard error for each electrode.

**Supplementary Figure 3:**
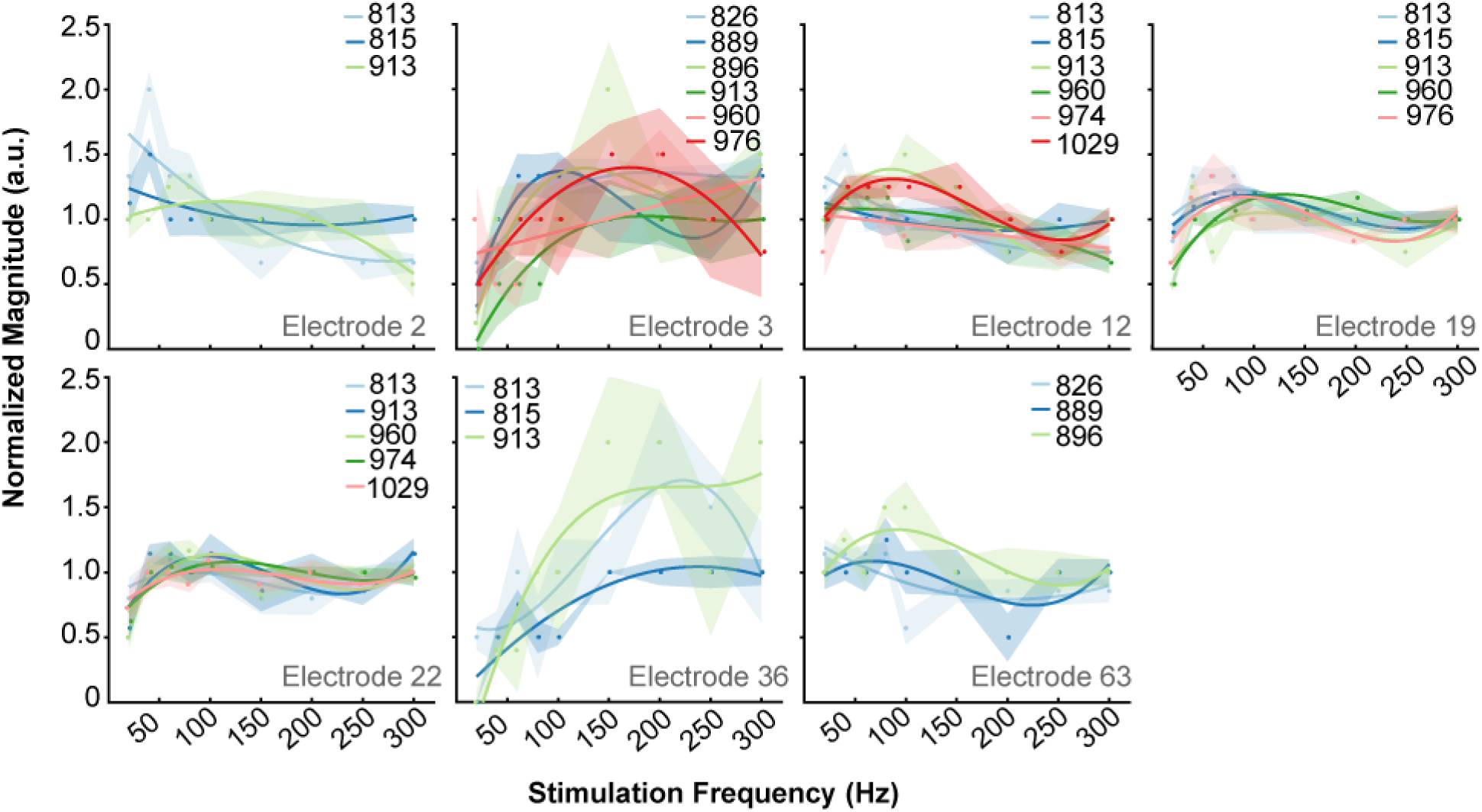
Electrodes maintain same frequency-intensity relationships over time. Plots of magnitude estimation results on all electrodes tested three or more times. Each set of points and the corresponding fit indicate a single post implant date, as indicated in the legend. Data from each test session were normalized by the median intensity. Different colors show different post-implant dates in each plot as indicated by the legend. Shaded regions show the smoothed standard error.

**Supplementary Figure 4:**
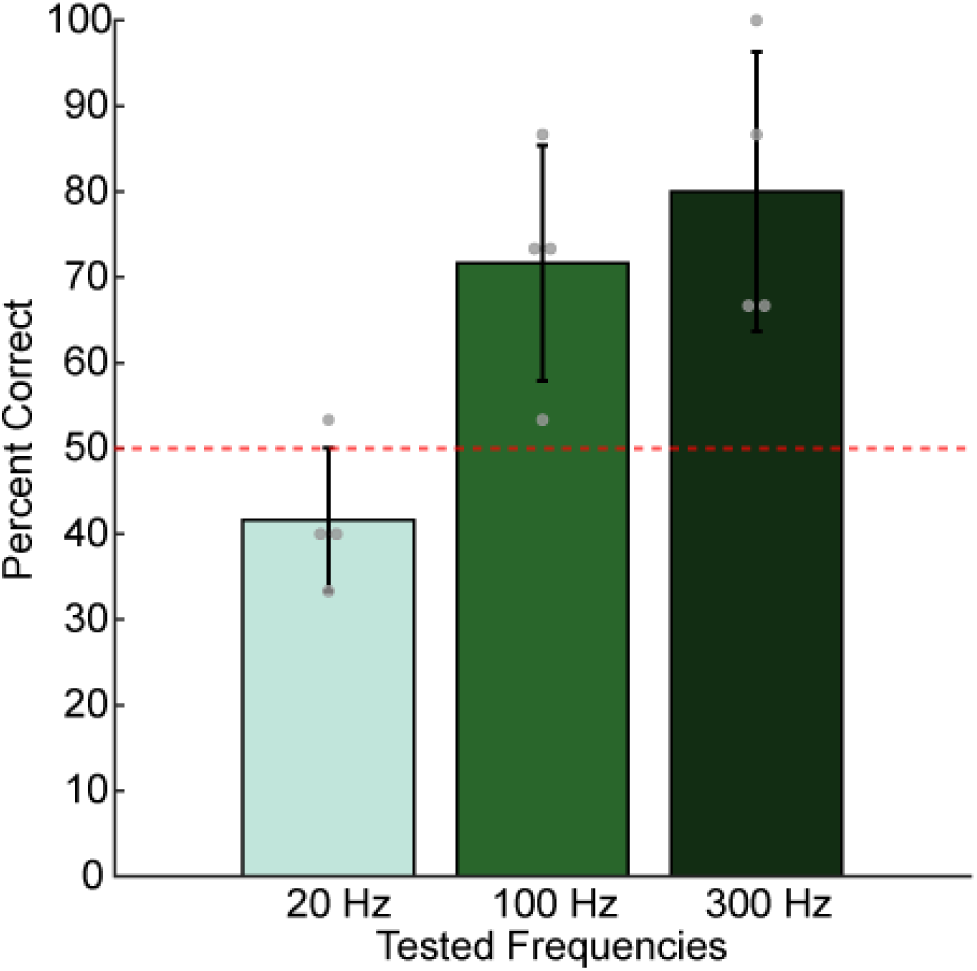
Higher pulse frequencies always improved detection at perithreshold current amplitudes. Bar plots showing the probability of correctly identifying the window containing a stimulus train with different pulse frequencies at a fixed perithreshold current amplitude (6-16 μA). Each bar represents the mean detection accuracy at each pulse frequency on four tested electrodes. The error bar indicates the standard deviation. The grey dots show the individual electrode performance accuracies. Chance performance was 50% and is indicated with the red dotted line.

**Supplementary Figure 5:**
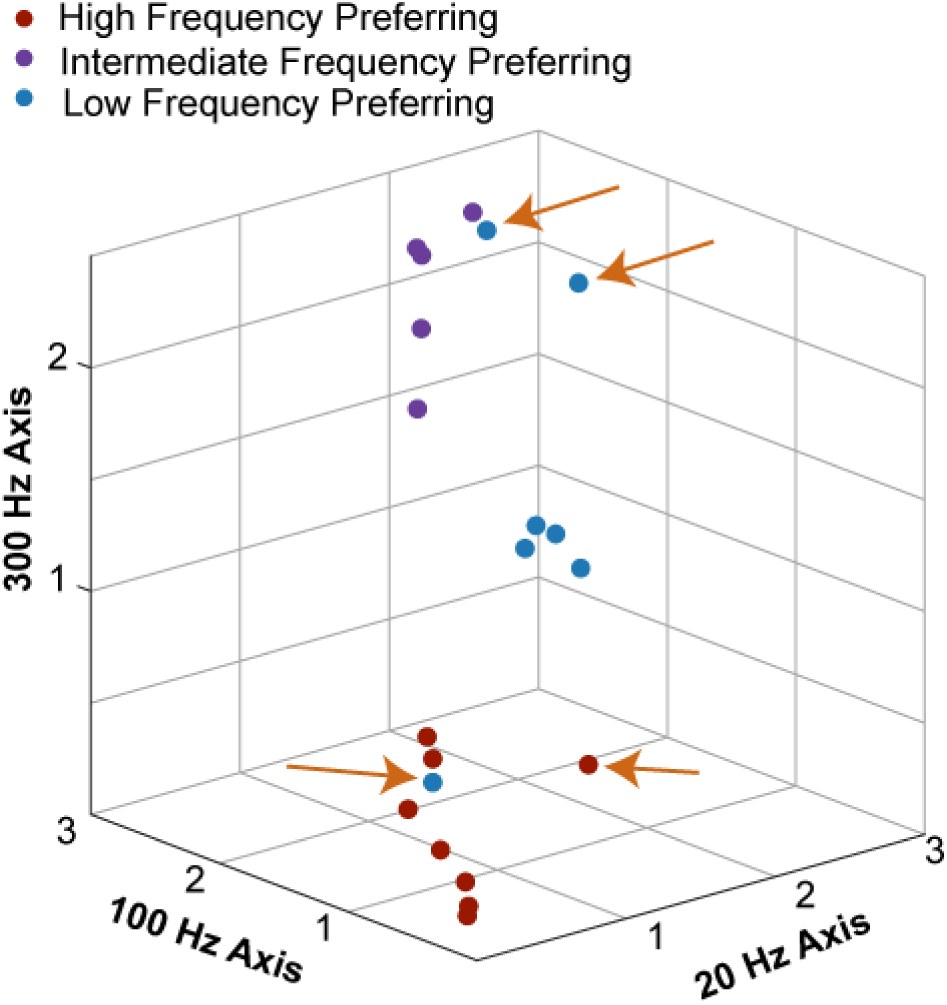
Clustering by evoked qualities results in nearly identical clusters to those identified from perceived intensity. Individual electrodes are plotted on the same axes as shown in Supplementary Figure 2A. The data were clustered in a ten-dimensional quality space and then plotted in the three-dimensional frequency-intensity space. Clusters are labeled based on the three categories defined by frequency-intensity responses. For example, the blue point in the lower portion of the figure represents an electrode that shared similar frequency-intensity properties with other high frequency preferring electrodes but shared qualities that were similar to low frequency preferring electrodes. However, the majority of the electrodes were identified as being in the same clusters regardless of whether the clustering was performed on quality or frequency-intensity data. Electrodes that were classified differently between quality and frequency-intensity data are indicated with orange arrows.

## References

Anselin, L., 1995. Local Indicators of Spatial Association—LISA. Geogr. Anal. 27, 93–115.

Armenta Salas, M., Bashford, L., Kellis, S., Jafari, M., Jo, H., Kramer, D., Shanfield, K., Pejsa, K., Lee, B., Liu, C.Y., Andersen, R.A., 2018. Proprioceptive and cutaneous sensations in humans elicited by intracortical microstimulation. Elife e32904.

Butovas, S., Schwarz, C., 2007. Detection psychophysics of intracortical microstimulation in rat primary somatosensory cortex. Eur. J. Neurosci. 25, 2161–2169.

Callier, T., Brantly, N.W., Caravelli, A., Bensmaia, S.J., 2020. The frequency of cortical microstimulation shapes artificial touch. Proc. Natl. Acad. Sci. U. S. A. 117, 1191–1200.

Chen, L.M., Friedman, R.M., Ramsden, B.M., LaMotte, R.H., Roe, A.W., 2001. Fine-scale organization of SI (area 3b) in the squirrel monkey revealed with intrinsic optical imaging. J. Neurophysiol. 86, 3011–3029.

Cogan, S.F., 2008. Neural Stimulation and Recording Electrodes. Annu. Rev. Biomed. Eng. 10, 275–309.

Collinger, J.L., Gaunt, R.A., Schwartz, A.B., 2018. Progress towards restoring upper limb movement and sensation through intracortical brain-computer interfaces. Curr. Opin. Biomed. Eng.

Dadarlat, M.C., O’Doherty, J.E., Sabes, P.N., 2014. A learning-based approach to artificial sensory feedback leads to optimal integration. Nat. Neurosci. 18, 138–144.

Dicarlo, J.J., Johnson, K.O., Hsiao, S.S., 1998. Structure of receptive fields in area 3b of primary somatosensory cortex in the alert monkey. J. Neurosci. 18, 2626–2645.

Fetz, E.E., 2015. Restoring motor function with bidirectional neural interfaces. In: Progress in Brain Research. Elsevier, pp. 241–252.

Fifer, M.S., McMullen, D.P., Thomas, T.M., Osborn, L.E., Nickl, R., Candrea, D., Pohlmeyer, E., Thompson, M., Anaya, M., Schellekens, W., Ramsey, N., Bensmia, S., Anderson, W., Wester, B., Crone, N., Celnik, P., Cantarero, G., Tenore, F. V, 2020. Intracortical microstimulation of human fingertip sensations. medRxiv.

Flesher, S.N., Collinger, J.L., Foldes, S.T., Weiss, J.M., Downey, J.E., Tyler-Kabara, E.C., Bensmaia, S.J., Schwartz, A.B., Boninger, M.L., Gaunt, R.A., 2016. Intracortical microstimulation of human somatosensory cortex. Sci. Transl. Med. 8, 1–11.

Flesher, S.N., Downey, J.E., Weiss, J.M., Hughes, C.L., Herrera, A.J., Tyler-Kabara, E.C., Boninger, M.L., Collinger, J.L., Gaunt, R.A., 2019. Restored tactile sensation improves neuroprosthetic arm control. bioRxiv 653428.

Fridman, G.Y., Blair, H.T., Blaisdell, A.P., Judy, J.W., 2010. Perceived intensity of somatosensory cortical electrical stimulation. Exp. Brain Res. 203, 499–515.

Friedman, R.M., Chen, L.M., Roe, A.W., 2004. Modality maps within primate somatosensory cortex. Proc. Natl. Acad. Sci. U. S. A. 101, 12724–9.

Godfrey, S.B., Bianchi, M., Bicchi, A., Santello, M., 2016. Influence of force feedback on grasp force modulation in prosthetic applications: A preliminary study. In: Proceedings of the Annual International Conference of the IEEE Engineering in Medicine and Biology Society, EMBS. pp. 5439–5442.

Graczyk, E.L., Schiefer, M.A., Saal, H.P., Delhaye, B.P., Bensmaia, S.J., Tyler, D.J., 2016. The neural basis of perceived intensity in natural and artificial touch. Sci. Transl. Med. 8, 362ra142 LP–362ra142.

Heming, E., Sanden, A., Kiss, Z.H.T., 2010. Designing a somatosensory neural prosthesis: percepts evoked by different patterns of thalamic stimulation. J. Neural Eng. 7, 064001.

Hollins, M., Roy, E.A., 1996. Perceived Intensity of Vibrotactile Stimuli: The Role of Mechanoreceptive Channels. Somatosens. Mot. Res. 13, 273–286.

Hughes, C.L., Herrera, A., Gaunt, R., Clinical, J.C.-H. of, 2020, U., 2020. Bidirectional brain-computer interfaces. Handb. Clin. Neurol. 168, 163–181.

Johansson, R.S., Flanagan, J.R., 2009. Coding and use of tactile signals from the fingertips in object manipulation tasks. Nat. Rev. Neurosci. 10, 345–59.

Kapfer, C., Glickfeld, L.L., Atallah, B. V., Scanziani, M., 2007. Supralinear increase of recurrent inhibition during sparse activity in the somatosensory cortex. Nat. Neurosci. 10, 743–753.

Kim, L.H., McLeod, R.S., Kiss, Z.H.T., 2018. A new psychometric questionnaire for reporting of somatosensory percepts. J. Neural Eng.

Kim, S., Callier, T., Tabot, G. a, Gaunt, R.A., Tenore, F. V, Bensmaia, S.J., 2015a. Behavioral assessment of sensitivity to intracortical microstimulation of primate somatosensory cortex. Proc. Natl. Acad. Sci. 112, 15202–15207.

Kim, S., Callier, T., Tabot, G.A., Tenore, F. V., Bensmaia, S.J., 2015b. Sensitivity to microstimulation of somatosensory cortex distributed over multiple electrodes. Front. Syst. Neurosci. 9, 47.

Large, A.M., Vogler, N.W., Canto-Bustos, M., Friason, F.K., Schick, P., Oswald, A.M.M., 2018. Differential inhibition of pyramidal cells and inhibitory interneurons along the rostrocaudal axis of anterior piriform cortex. Proc. Natl. Acad. Sci. U. S. A. 115, E8067–E8076A.

Leek, M.R., 2001. Adaptive procedures in psychophysical research. Percept. Psychophys. 63, 1279–1292.

Levitt, H., 1971. Transformed Up-Down Methods in Psychoacoustics. J. Acoust. Soc. Am. 49, 467–477.

Luna, V.M., Pettit, D.L., 2010. Asymmetric rostro-caudal inhibition in the primary olfactory cortex. Nat. Neurosci. 13, 533–535.

Michelson, N.J., Eles, J.R., Vazquez, A.L., Ludwig, K.A., Kozai, T.D.Y., 2019. Calcium activation of cortical neurons by continuous electrical stimulation: Frequency dependence, temporal fidelity, and activation density. J. Neurosci. Res. 97, 620–638.

Mountcastle, V.B., Talbot, W.H., Sakata, H., Hyvärinen, J., 1969. Cortical neuronal mechanisms in flutter-vibration studied in unanesthetized monkeys. Neuronal periodicity and frequency discrimination. J. Neurophysiol. 32, 452–484.

Muniak, M.A., Ray, S., Hsiao, S.S., Dammann, J.F., Bensmaia, S.J., 2007. The Neural Coding of Stimulus Intensity: Linking the Population Response of Mechanoreceptive Afferents with Psychophysical Behavior. J. Neurosci. 27, 11687–11699.

Nowak, D.A., Glasauer, S., Hermsdörfer, J., 2004. How predictive is grip force control in the complete absence of somatosensory feedback? Brain 127, 182–192.

Nowak, D.A., Glasauer, S., Hermsdörfer, J., 2013. Force control in object manipulation-A model for the study of sensorimotor control strategies. Neurosci. Biobehav. Rev. 37, 1578–1586.

Nowak, D.A., Hermsdörfer, J., 2006. Predictive and reactive control of grasping forces: On the role of the basal ganglia and sensory feedback. Exp. Brain Res. 173, 650–660.

Pei, Y.C., Denchev, P. V., Hsiao, S.S., Craig, J.C., Bensmaia, S.J., 2009. Convergence of submodality-specific input onto neurons in primary somatosensory cortex. J. Neurophysiol. 102, 1843–1853.

Penfield, W., Boldrey, E., 1937. Somatic motor and sensory representation in the cerebral cortex of man as studied by electrical stimulation. Brain 60, 389–443.

Reed, J.L., Qi, H.X., Zhou, Z., Bernard, M.R., Burish, M.J., Bonds, A.B., Kaas, J.H., 2010. Response properties of neurons in primary somatosensory cortex of owl monkeys reflect widespread spatiotemporal integration. J. Neurophysiol. 103, 2139–2157.

Romo, R., Hernández, A., Zainos, A., Brody, C.D., Lemus, L., 2000. Sensing without touching: psychophysical performance based on cortical microstimulation. Neuron 26, 273–8.

Romo, R., Hernández, A., Zainos, A., Salinas, E., 1998. Somatosensory discrimination based on cortical microstimulation. Nature 392, 387–390.

Rosenbaum, R., Rubin, J., Doiron, B., 2012. Short term synaptic depression imposes a frequency dependent filter on synaptic information transfer. PLoS Comput. Biol. 8.

Saal, H.P., Bensmaia, S.J., 2014. Touch is a team effort: Interplay of submodalities in cutaneous sensibility. Trends Neurosci. 37, 689–697.

Schmidt, E.M., Bak, M.J., Hambrecht, F.T., Kufta, C. V., O’Rourke, D.K., Vallabhanath, P., 1996. Feasibility of a visual prosthesis for the blind based on intracortical micro stimulation of the visual cortex. Brain 119, 507–522.

Semprini, M., Bennicelli, L., Vato, A., 2012. A parametric study of intracortical microstimulation in behaving rats for the development of artificial sensory channels. In: Proceedings of the Annual International Conference of the IEEE Engineering in Medicine and Biology Society, EMBS. pp. 799–802.

Silberberg, G., Markram, H., 2007. Disynaptic Inhibition between Neocortical Pyramidal Cells Mediated by Martinotti Cells. Neuron 53, 735–746.

Sur, M., Wall, J.T., Kaas, J.H., 1981. Modular segregation of functional cell classes within the postcentral somatosensory cortex of monkeys. Science (80-.). 212, 1059–1061.

Sur, M., Wall, J.T., Kaas, J.H., 1984. Modular distribution of neurons with slowly adapting and rapidly adapting responses in area 3b somatosensory cortex in monkeys. J. Neurophysiol. 51, 724–744.

Tsodyks, M. V., Markram, H., 1997. The neural code between neocortical pyramidal neurons depends on neurotransmitter release probability. Proc. Natl. Acad. Sci. U. S. A. 94, 719–723.

Verrillo, R.T., Fraioli, A.J., Smith, R.L., 1969. Sensation magnitude of vibrotactile stimuli. Percept. Psychophys. 6, 366–372.

